# Phylogenetic Analysis of TET2 Gene Variants in Pakistani Acute Myeloid Leukemia Patients

**DOI:** 10.1101/2025.09.03.674098

**Authors:** Mawada Elmagboul Abdalla Abakar, Mehad Almagboul Abdalla Abaker

## Abstract

**Background:** Acute myeloid leukemia (AML) is a heterogeneous malignancy caused by the proliferation of neoplastic myeloid progenitor cells. Mutations in ten-eleven translocation methylcytosine dioxygenase 2 (TET2), a key regulator of DNA demethylation, are common in AML and play a role in its progression and response to treatment.

**Methods:** We performed a phylogenetic analysis of 25 TET2 gene sequences (12 AML and 13 normal) from Pakistani individuals using multiple sequence alignment, phylogenetic tree construction, and distance matrix evaluation. We also identified population-specific expression patterns and mutational hotspots.

**Results:** AML samples showed significant deletions in the catalytic domain (positions 693– 876) and lower sequence conservation compared to controls. These deletions may impair TET2 enzymatic activity and disrupt the epigenetic regulation. The Phylogenetic tree also showed two significant clades supported by high bootstrap values, differentiating AML from control individuals in the evolutionary direction.

**Conclusion:** Our findings reveal distinct TET2 mutational hotspots associated with acute myeloid leukemia (AML) in the Pakistani population. These results have implications for targeted epigenetic therapy and motivate large-scale, cross-population research to unravel the global effects of TET2 variations.

## 1. Introduction

Acute myeloid leukemia (AML) is an aggressive cancer affecting hematopoietic progenitor cells, characterized by uncontrolled proliferation, abnormal differentiation of myeloid precursor cells, complex clonal structure, and disease progression driven by various genetic and epigenetic alterations[1], [2], [3], [4]. TET2 (ten-eleven translocation 2) is a member of the TET family that encodes DNA dioxygenases, which regulate DNA demethylation and other epigenetic modifications. TET2 functions as a key enzyme that plays a critical role in converting 5-methylcytosine (5mC) to 5-hydroxymethylcytosine (5hmC), an essential step in DNA demethylation[5]. TET2, a key regulator of DNA methylation dynamics, has emerged as a critical player in hematopoietic development and leukemogenesis. Normal hematopoiesis is a highly regulated process in which hematopoietic stem and progenitor cells (HSPCs) differentiate into various mature blood cell lineages. TET2 plays a crucial role in this process by ensuring proper gene expression and maintaining the balance between self-renewal and differentiation of HSPCs [6].

Recent studies have reported significant differences in TET2 expression patterns between different human populations, indicating the presence of complex molecular population-specific mechanisms in AML pathogenesis [1], [7], [8]. Recently TET2 Mutations have been found in 7-28% of AML cases, more often in elderly individuals, and therefore are a key requirement in leukemogenesis. These mutations can cause the loss of enzymatic activity, thus, causing epigenetic dysregulation and altering the growth and differentiation potential of hematopoietic stem and progenitor cells (HSPCs) [1], [9]. Dysregulation of TET2 function has the potential to affect the epigenetic regulation of HSPCs, leading to inappropriate oncogene activation and inhibition of tumor suppressor genes, eventually leading to AML development [10], [11].

### 1.1 TET2 Gene Function and Structure

The TET2 gene is located on chromosome 4q24, has an approximate range of 133.9 Kb, and the full-length gene product is 2,002 amino acids. This gene is expressed ubiquitously throughout the hematopoietic system, with a high abundance in subsets of HSPCs and mature myeloid and lymphoid cells. It converts 5-methylcytosine (5mC) to 5-formylcytosine (5fC), 5-hydroxymethylcytosine (5hmC), and 5-carboxycytosine (5caC), which is an essential step in active DNA demethylation and maintenance of normal hematopoiesis. [12], [13], [14], [15].

### 1.2 TET2 Mutations in Clonal Hematopoiesis and AML

Mutations in the TET2 gene are not only present in AML but are also found in healthy elderly people, leading to clonal hematopoiesis (CH). Clonal hematopoiesis of indeterminate potential (CHIP) is a subset of CH involving clonal expansion of somatic blood cell clones with mutations in the driver genes of leukemogenesis, with TET2 being one of the most commonly affected genes[16]. CHIP is distinguished by a higher risk of hematological malignancies, including AML, although not all individuals with CHIP develop leukemia[17], [18]. TET2 mutations are frequently observed in CHIP, highlighting their role in the pathogenesis of early myeloid malignancies. TET2 mutations often occur early in the development of acute myeloid leukemia (AML) and are strongly correlated with age, particularly in older individuals[19]. The role of TET2 mutations in AML highlights their importance in understanding the disease. These mutations contribute to cell growth and clonal evolution, which drive the progression of leukemia. Mutations that lead to loss of function in TET2 are common early events during clonal hematopoiesis. They act as pre-leukemic “founder” lesions that persist during chemotherapy and influence subsequent evolutionary trajectories. TET2 encodes an Fe(II)-α-KG-dependent dioxygenase. This enzyme converts 5-methylcytosine into 5-hydroxymethylcytosine (5-hmC). This function is crucial for controlling the gene expression and cellular processes. When TET2 loses its activity, the levels of 5-hmC decrease. This change disrupts DNA demethylation and leads to the dysregulation of lineage-specific transcription programs that encourage self-renewal and block terminal myeloid differentiation. In AML, TET2 mutations often occur alongside other mutations such as CBL and SRSF2 in TET2-mutant cases. In contrast, DNMT3A-mutant cases tended to favor NPM1/FLT3, resulting in different evolutionary paths and clinical behaviors. Importantly, truncating (frameshift/nonsense) or biallelic TET2 alterations are associated with worse outcomes and diminished efficacy of allogeneic transplantation in AML patients. TET2 mutations create an epigenetic and clonal foundation that facilitates the acquisition of secondary driver lesions such as signalling or transcription factor mutations. This accelerates the progression of clonal hematopoiesis to AML.[1], [2], [20], [21], [22].

### 1.3 Clinical Significance and Prognostic Impact

In AML, TET2 mutations have important interactions with other mutations, various prognostic effects, and offer new treatment possibilities. TET2 mutations exhibit different co-mutation patterns that affect clinical outcomes. For example, AML patients with both TET2 and NPM1 mutations are often seen in AML cases and have intermediate prognostic significance[1], [23], [24]. In contrast, TET2 mutations show mutual exclusivity with IDH1/2 mutations, while they commonly co-occurring with other epigenetic regulators, including ASXL1, SRSF2, and DNMT3A, although these co-mutation patterns do not uniformly affect prognosis[1], [2], [22], [25], [26], [27], [28].

The significance of TET2 mutations in patient outcomes is not uniform and varies depending on the clinical setting. While most large AML cohorts demonstrate no significant differences in overall survival, event-free survival, or relapse rates among TET2-mutated and wild-type patients, specific subgroups showed more pronounced effects[22], [29], [30].TET2 mutations are preferentially enriched within cytogenetically normal AML (CN-AML) and intermediate-risk cytogenetic categories, occuring in 18-23% of cases [31]. Within these subgroups, TET2 mutations may be correlated with decreased overall survival and reduced event-free survival, particularly when present at high variant allele frequencies, although these associations lack consistency across different cohorts [9], [22], [29], [30], [32], [33], [34]. Recent studies have suggested that the prognostic impact may be more pronounced in older patients and those with specific co-mutation profiles[9], [23], [35].

Clinically, AML patients harboring TET2 mutations present with distinctive laboratory features, including significantly elevated white blood cell (WBCs) counts at diagnosis. Additionally, some groups show reduced platelet counts and increased blast percentages[1], [36]. These mutations are predominantly observed in patients with normal karyotypes, in addition to, exhibit a strong correlation with intermediate-risk cytogenetic profiles[1], [34]. The biological basis for these clinical associations is relates to TET2’s role in hematopoietic differentiation, where loss-of-function mutations confer a competitive advantage to mutant clones through enhanced self-renewal capacity, altered differentiation programs, and potential resistance to conventional therapeutic approaches[36], [37], [38].

The therapeutic implications of TET2 alterations represent an area of active clinical investigation and emerging potential. TET2-mutated AML cells demonstrated enhanced sensitivity to hypomethylating therapies, such as 5-azacytidine and decitabine, in both myelodysplastic syndrome and AML patient populations[39], [40], [41]. This sensitivity appears to result from the inability of TET2-deficient cells to properly regulate DNA methylation patterns, making them more dependent on DNA methyltransferase activity and, thus, more susceptible to hypomethylating agent-induced cytotoxicity[42], [43], [44], [45]. Additionally, vitamin C supplementation has shown therapeutic potential by enhancing the residual activity of functional TET enzymes and partially restoring normal DNA demethylation patterns in cells with TET2-mutations[1], [46]. Emerging targeted approaches include specific TET2 inhibitors and strategies combined with PARP inhibitors based on the synthetic lethal relationship between TET2 deficiency and DNA repair pathway dependency[47], [48].

### 1.4 Phylogenetic Analysis in Cancer Research

Understanding the evolutionary relationships of TET2 in various populations may provide insight into the selective pressures underlying leukemogenesis. Phylogenetic methods can be used to classify AML samples based on their gene expression profiles. These samples can grouped into subtypes based on their maturation status. Such analyses can potentially reveal population-specific expression patterns of TET2. Phylogenetic analysis can also help rank cancer subtypes according to their similarity to stem cells, with poorly differentiated cancers having gene expression patterns more similar to those of stem cells. This approach is based on the concept that cancers may arise from an interruption in the differentiation process[33], [49], [50].

### 1.5 Study Objectives

However, details of the evolutionary process of TET2 in AML have not yet been fully established. This study aimed to elucidate the evolutionary conservation and divergence between population-specific TET2 expression patterns and AML pathogenesis through comprehensive phylogenetic analysis. By examining the evolutionary relationships and molecular signatures of TET2 variants across the Pakistani population, we aimed to uncover population-specific expression patterns of TET2 and contribute to a deeper understanding of the mechanisms underlying AML progression, thereby facilitating the advancement of targeted therapeutic strategies.

## 2. Material and Method

### 2.1 Gene Sequence Retrieval and Data Preparation

We obtained 25 DNA sequences of the TET2 gene from acute myeloid leukemia(AML) patients and normal controls (13 normal and 12 AML patients) from Pakistan using the publicly available database of the National Center for Biotechnology Information (NCBI) (https://www.ncbi.nlm.nih.gov). The sequence data were accessed for research purposes on August 10, 2020, and December 19, 2020. All sequences were previously deposited in the NCBI database as de-identified, publicly available data. The authors did not have access to any information that could identify individual participants during or after data collection, as all sequences were anonymized prior to public deposition in accordance with NCBI data sharing policies. These sequences were used to study the evolution and phylogenetic relationship of TET2 in AML patients (Table 1).

**Table 1:**
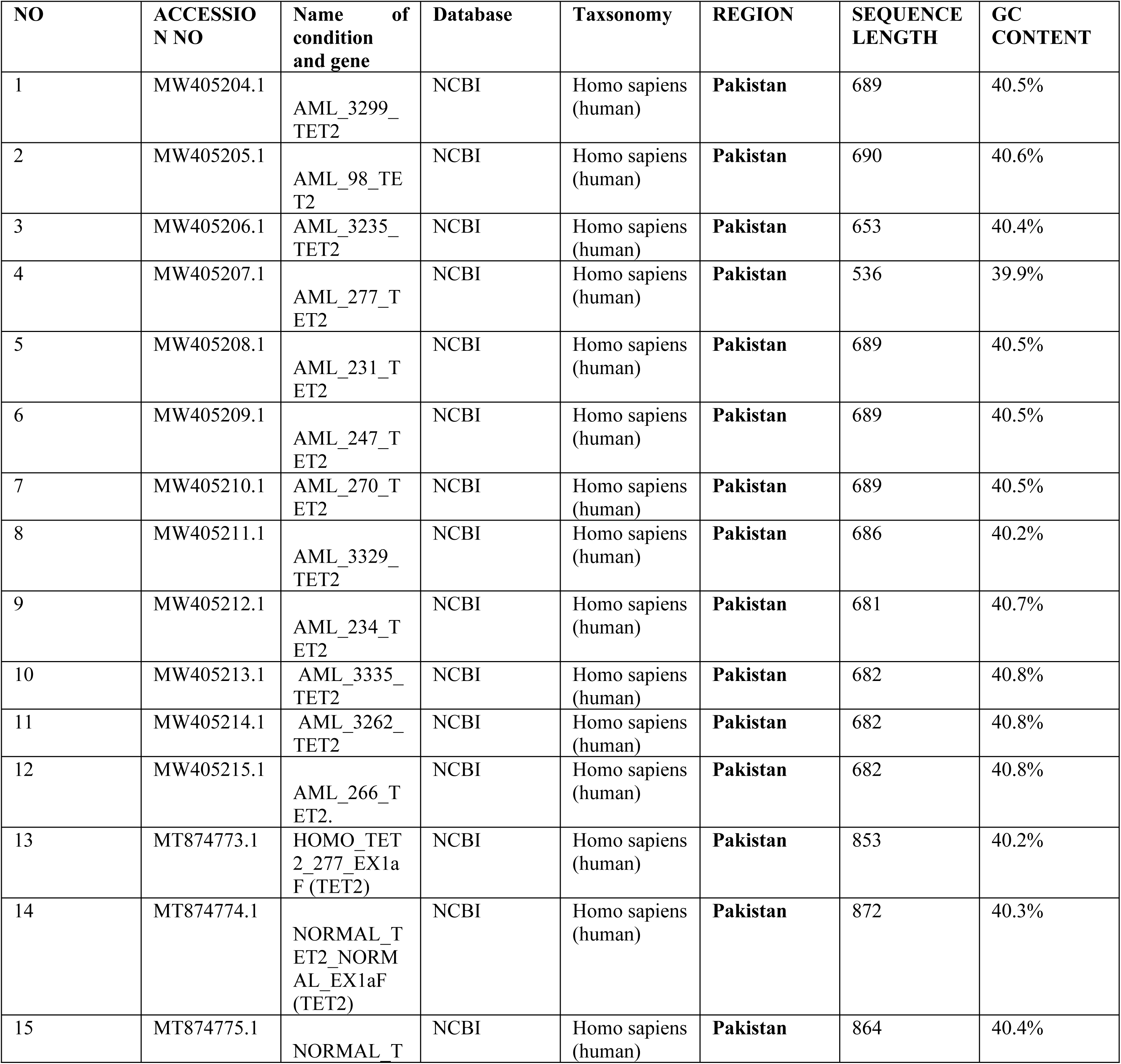
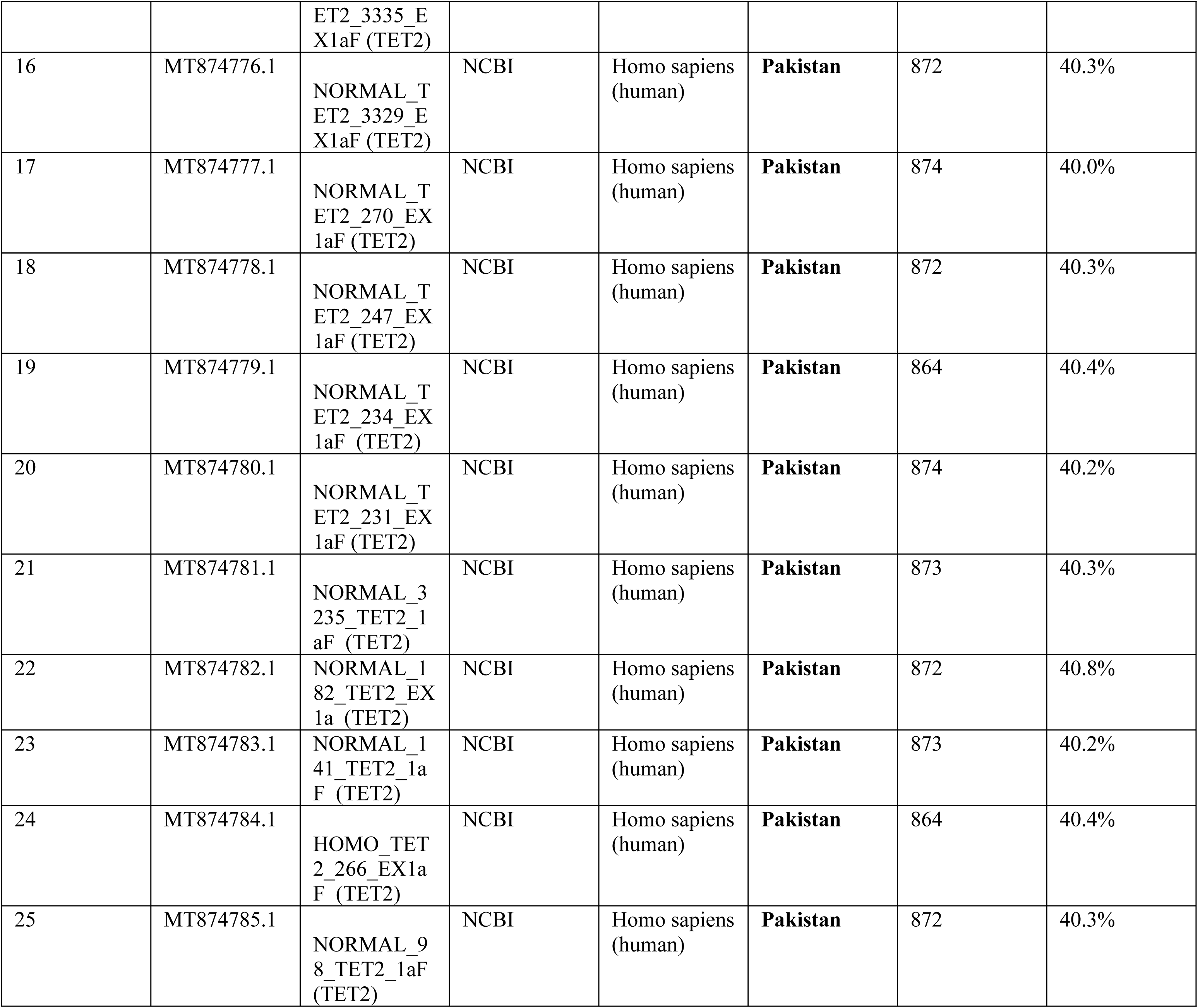
Characteristics of the 25 TET2 gene sequences from AML patients and normal controls used in this Study.

### 2.2 Multiple Sequence Alignment

To conduct multiple sequence alignment (MSA), the retrieved TET2 gene sequences were uploaded to Geneious software, which provides a user-friendly interface with a comprehensive range of tools for sequence analysis, including sequence alignment. The Geneious Alignment tool in Geneious software v9.1.8 was utilized due to its speed and accuracy during the alignment process. The alignment process aimed to identify conserved regions and align the sequences based on shared similarities and homologies. Gaps and mismatches were introduced to ensure proper sequence alignment.

After alignment was completed, an output file was obtained, which included the aligned sequences and corresponding gap positions. This alignment file was used for further analyses, including phylogenetic tree construction and determination of conserved regions.

### 2.3 Phylogenetic Tree Construction

Following sequence alignment, we conducted a phylogenetic analysis using the Tamura-Nei genetic distance model and neighbor-joining method. This analysis utilized a bootstrap value of 1000 with 100 replicates to elucidate evolutionary relationships. A phylogenetic tree was constructed and edited using Geneious Tree Builder version 9.1.8 to elucidate evolutionary relatedness and divergence patterns.

A comparative analysis employing a distance matrix of the tree was conducted using Geneious software (https://www.geneious.com) based on statistical analysis to ascertain the positions of significant differences among the samples. The resulting alignment was used to identify conserved regions, insertions, deletions, and sequence variations in the genes. Conserved regions denote the critical functional elements of genes, whereas variations may indicate potential differences in disease susceptibility and progression.

### 2.4 Statistical Analysis

For comparative analysis and to determine the position of crucial importance among sequences, a distance matrix of the constructed phylogenetic tree was generated using Geneious v9.1.8, along with other statistical analyses, including Excel-based models. A distance matrix was used to identify divergence among the analyzed samples, whereas neighbor-joining clustering was used to assess the genetic distance and evolutionary relationship between sequences

## 3. Results and discussion

### 3.1 Comparative Sequence Analysis

The sequence data files selected from Pakistan exhibited various sequence lengths and GC contents, as shown in **Table 1**. Sequences with similar lengths and GC content indicate a shared evolutionary origin. In addition, sequences MW405213.1, MW405214.1, MW405215.1, and MT874782.1 showed a high GC content of 40.8%, 40.8%, 40.8%, and 40.8%. This suggests greater stability, which may influence the regulation of gene expression.

Interestingly, AML sequences exhibited greater variation in GC content than normal samples. This suggests that mutational events contribute to the development of leukemia. This observation aligns with recent findings that TET2 mutations are distributed throughout the gene, predominantly affecting the largest exons 3 and 11, with frameshift and nonsense mutations, leading to protein truncation and consequent loss of TET2 dioxygenase function [1], [11].

### 3.2 Multiple Sequence Alignment(MSA)

To investigate evolutionary relationships and their effects on AML resistance and progression in AML patients, multiple sequence alignments were performed on the retrieved gene sequences (Fig.1A). Multiple sequence alignment of 25 TET2 gene sequences from AML patients and normal controls revealed an overall alignment length of 876 nucleotides, with 787 identical sites, showing 89.8% identity between aligned sequences and a pairwise identity of 99.0%.

The statistical representation of the 25 ungapped aligned sequences included a mean length of 773.9 nucleotides, with a standard deviation of 103.4. The minimum sequence length was 536 nucleotides and the maximum sequence length was 874 nucleotides. Percentage base pair frequency distributions were **A**: 33.6% (total 6,504), **C**: 19.2 % (total 3,719), **G**: 21.0% (total 4,067), and **T**: 25.8% (total 4,993). The total GC content of the alignment was 40.4% (7,796).

Fig.1(a-c) shows the alignment representation of the sequence data, which indicates the similarities and differences occurring in different regions and locations within the sequences. Such variations raise the possibility of mutations, which may result in substitution, deletion, insertion, or translocation of nucleic acid sequences. These mutations are common throughout nucleotide sequences.

**Figure 1A:**
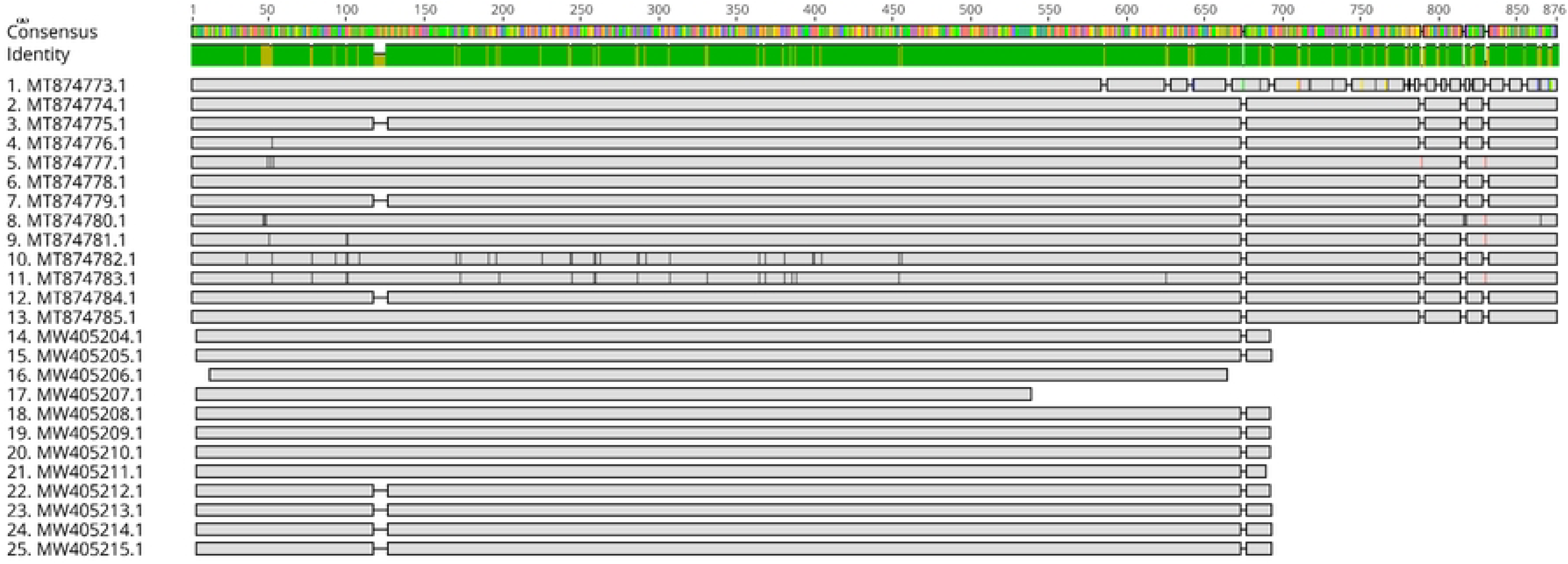
The alignment view of sequences (ranging 1-876 nucleotides) shows consensus similarities and variants among TET2 from AML patients and normal individuals, with the consensus identity view of the TET2 gene showing 55-65% similarity. Each row represents a TET2 sequence with nucleotide positions aligned across the samples. Accession numbers starting with (MW) are AML patients, and accession numbers starting with (MT) are normal. The top green bar (“Consensus Identity”) indicates regions of high sequence conservation (green = high identity, colored ticks = polymorphisms). Horizontal bars for each sample show the extent of sequence coverage; gaps indicate deletions or missing data. AML sequences displayed large gaps in the 693–876 region, whereas normal sequences were mostly intact.

**Figure 1B:**
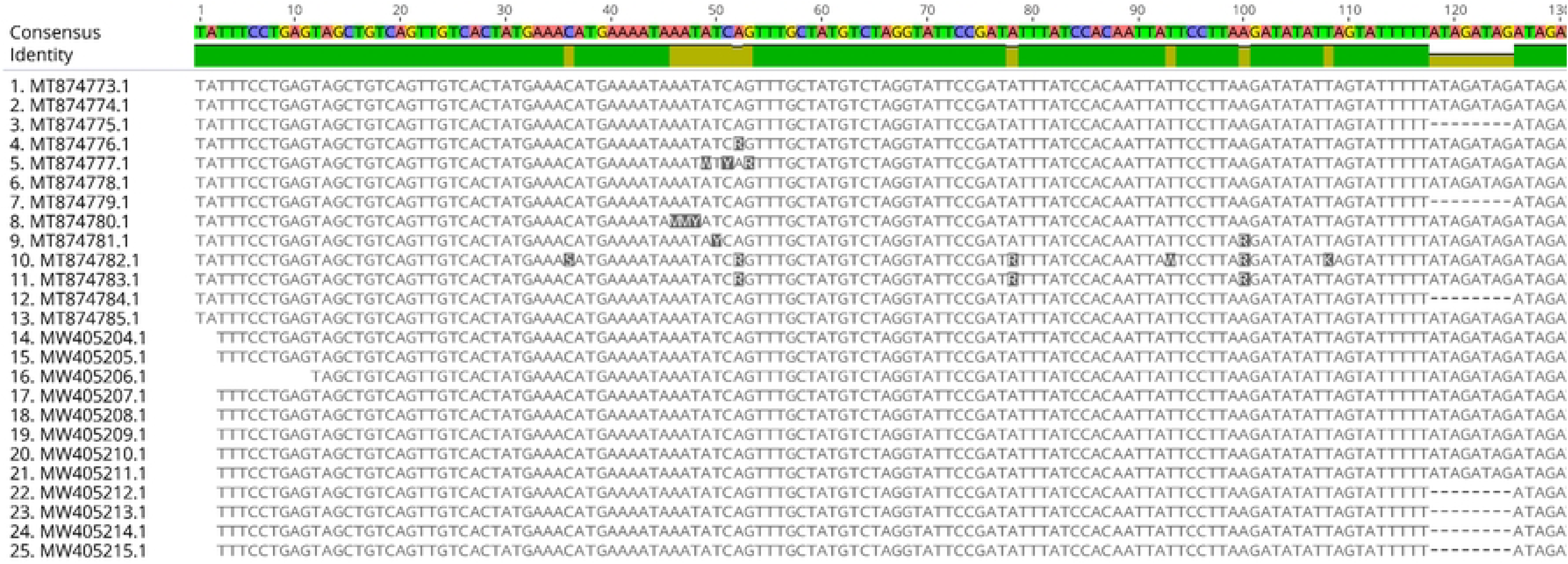
Sequence alignment view (nucleotide regions 1-130). The top green bar (“Consensus Identity”) indicates regions of high sequence conservation (green = high identity, colored ticks = polymorphisms). The upper part of the alignment shows high sequence conservation among most samples, with some scattered point mutations.

**Figure 1C:**
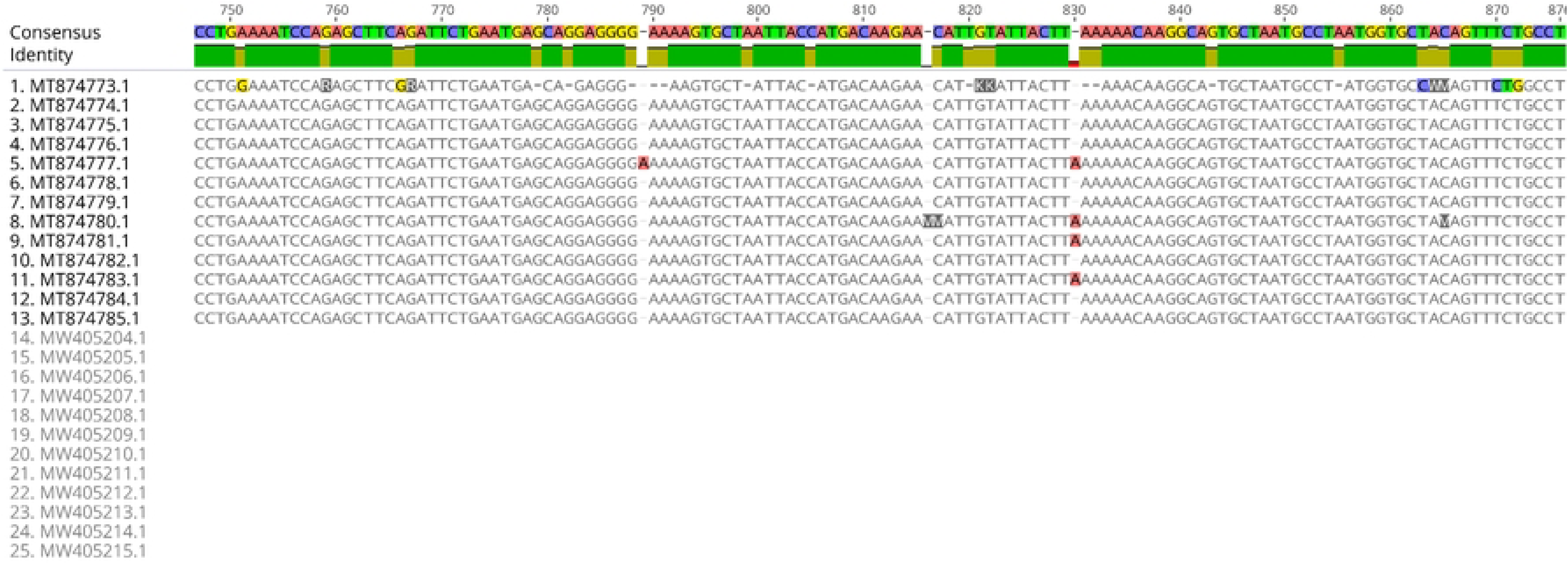
Sequence alignment view (nucleotide regions 750-876). The top green bar (“Consensus Identity”) indicates regions of high sequence conservation (green = high identity, colored ticks = polymorphisms). The lower part reveals extensive deletions in the AML sequences (MW), whereas the normal sequences (MT) remain largely intact.

The alignment view of the sequences (Fig.1A) is useful for quickly identifying conserved versus variable regions and visually correlating them with the disease status. All sequences from TET2 AML patients showed limited similarity and extensive variations compared to each other, with extensive deletions in the region from positions 693 to 876 (Fig.1A) compared to TET2 normal samples. This region is essential for TET2 catalytic activity, underscoring its potential role in TET2 dysfunction, leukemogenesis, and drug resistance.

These structural losses likely impair 5-hydroxymethylcytosine (5hmC) generation, echoing the reports of TET2 dysfunction in aggressive AML[19]. Deletions may result in the loss of genes responsible for the synthesis or formation of essential proteins or critical cellular structures. Consequently, such deletions may result in either a decrease or an increase in drug resistance.

The differences observed in this study may reflect the genetic diversity of TET2 among patients with AML and healthy controls. Fig(1b-1c) illustrate, with color coding, the regions of mutated genes within the sequences. The TET2 sample from AML patients clearly showed regions of nucleotide deletions that appeared as densely packed mutated areas. This may be associated with increased clinical virulence and patterns of drug resistance.

The AML-derived *TET2* sequences exhibited pronounced deletions in regions 693–876 (Fig. 1a,1c), which are specific to AML. This domain is critical for TET2 catalytic activity[14]. These structural losses likely impair 5hmC generation, highlighting the mutational hotspots that may drive disease progression and resistance to therapy.

#### Multiple Sequence Alignment Text Format

Sequences are typed in a one-letter code format, identifying nucleotides A, T, C, and G. Multiple sequences can be aligned and shown in this form with the identification of each sequence at the beginning. The alignment text format shown in (Fig.2) highlights the nucleotides containing deletions and variations in the base sequence between position 1 and more than 60 nucleotides. This shows the genetic diversity and mutation hotspots in the dataset. The variations in the sequences had varying features that might be clinically significant. In summary, as shown in the multiple sequence alignment text format (Fig.2), the AML samples had many deletions and hotspots of mutation, which might explain their potential role in TET2 dysfunction and AML pathogenesis.

**Figure 2:**
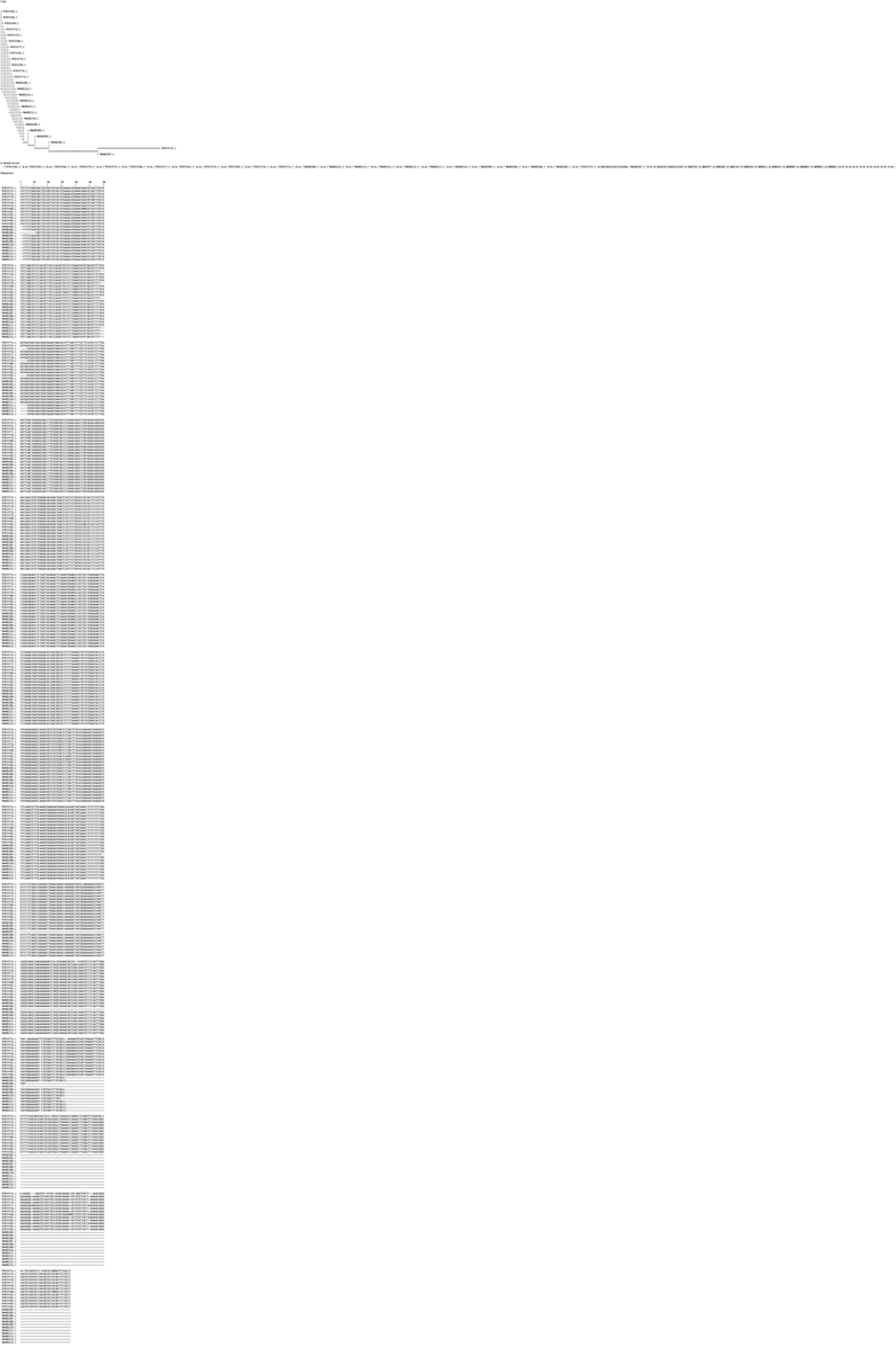
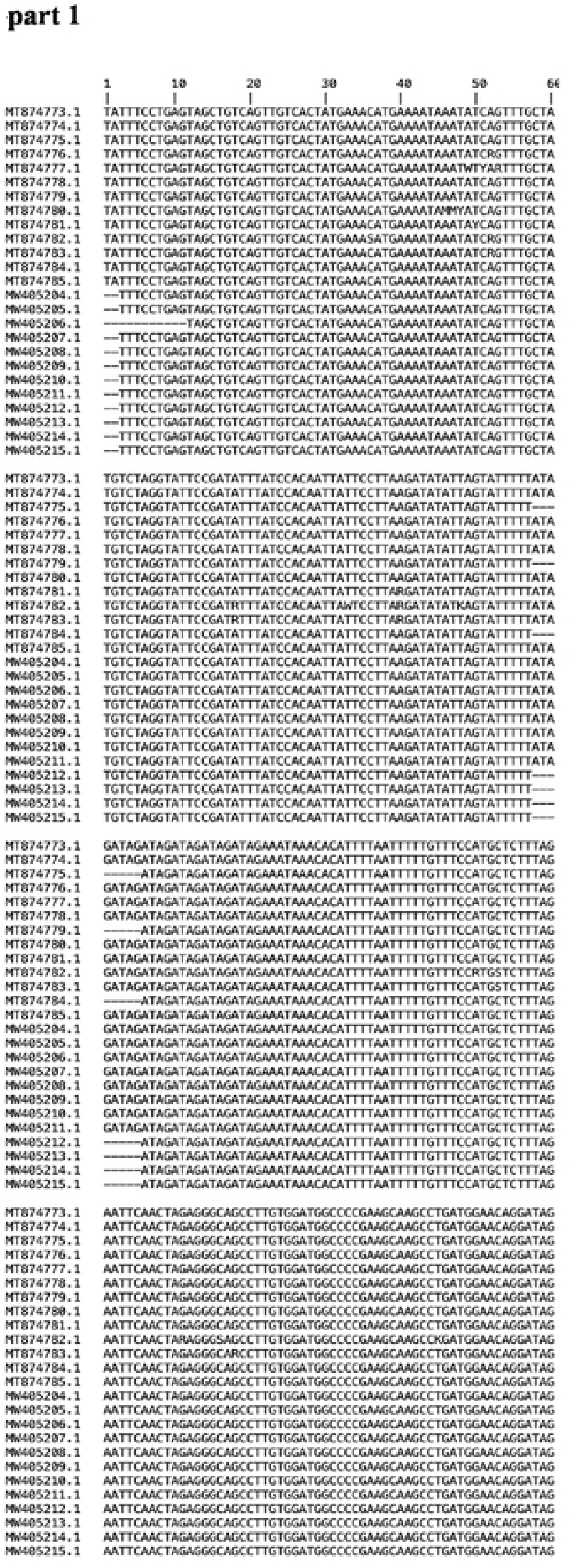
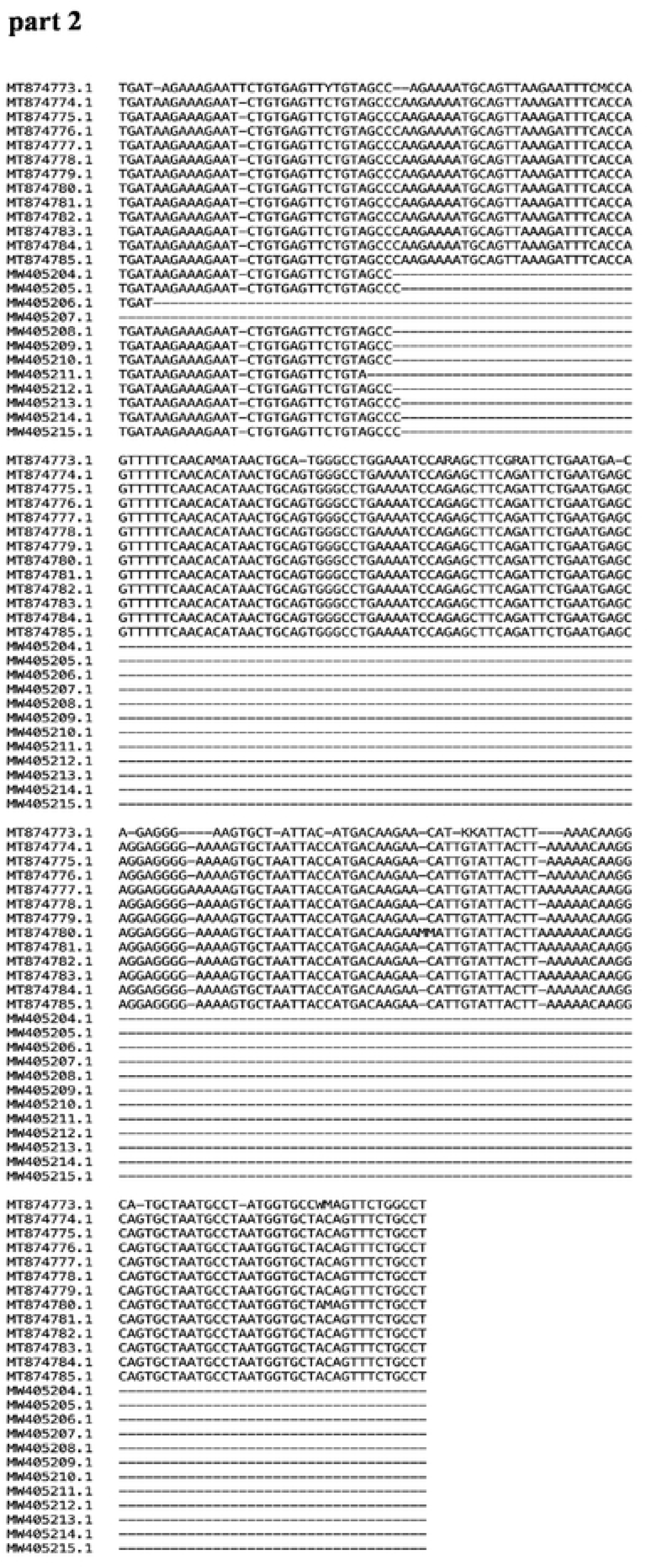
Sequence alignment in text format. Each row represents a TET2 sequence with nucleotide positions aligned across the samples. Gaps (represented by “–”) indicate deletions in the sequence.

### 3.3 Phylogenetic Analysis and Evolutionary Relationships

Phylogenetic analysis revealed different clustering patterns between AML and normal TET2 sequences, suggesting evolutionary differences associated with the disease status. The phylogenetic tree indicated the distinct separation of normal and AML-derived sample clusters with high bootstrap values supporting each branch (Fig.3). This provides strong statistical support that genetic divergence between normal and AML samples is strong and unlikely to be a random effect of sampling error, which identifies the robustness and clinical and evolutionary relevance of the observed sequence variation. This pattern highlights the evolutionary divergence and clonal growth-related nature of the AML pathogenesis.

**Figure 3:**
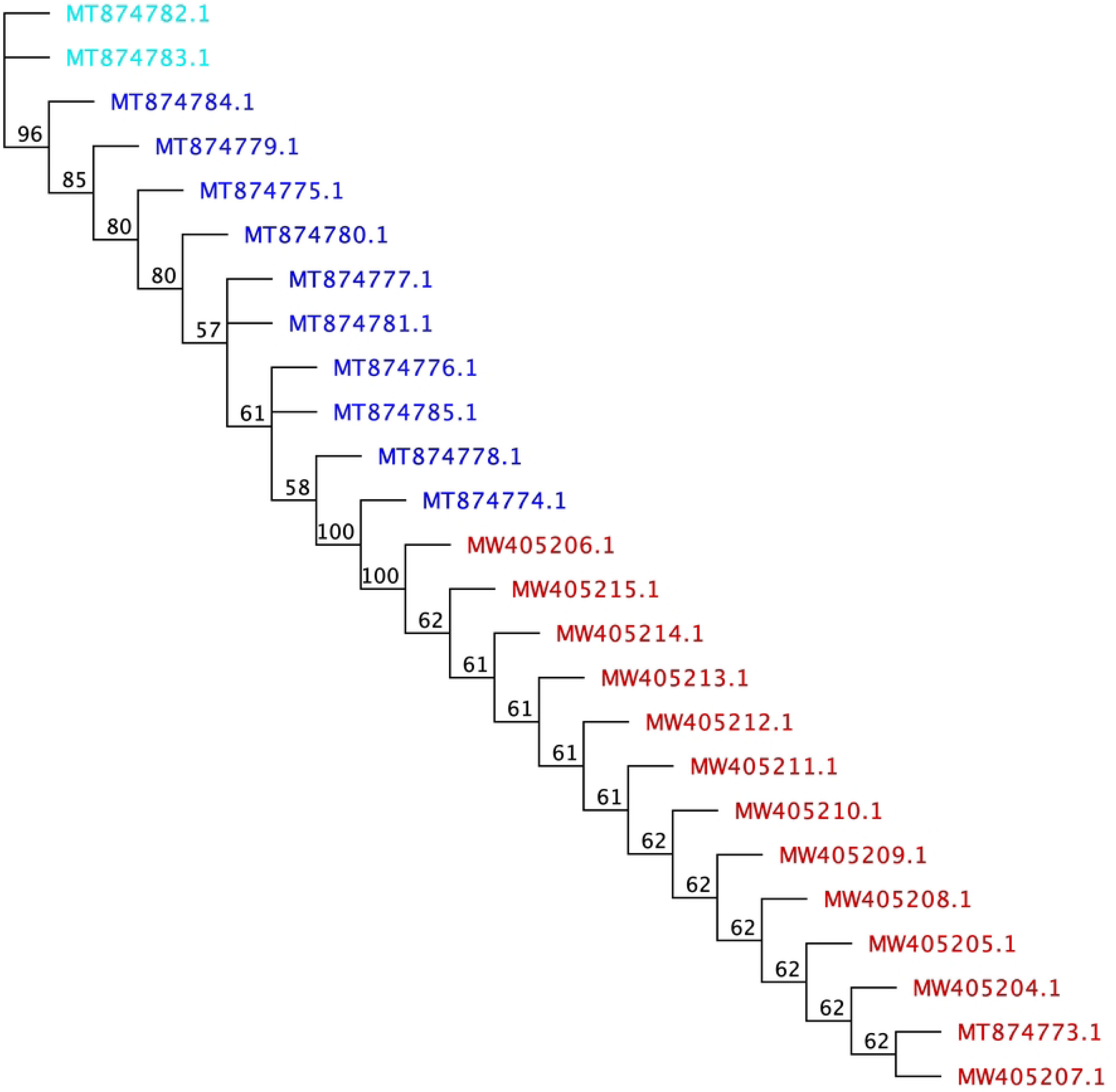
Phylogenetic Tree illustrating the relationships among various TET2 gene sequences related to AML patients and normal samples from Pakistan. Accession numbers starting with (MW) are AML patients, and accession numbers starting with (MT) are normal. A phylogenetic tree was generated using the neighbor-joining method.

The phylogenetic tree had 20 branches that emanating from the ancestral root, demonstrating the genetic diversity of the sequences. Each branch reflects a divergence in the parent, which can be of varying values as depicted by the bootstrap values. The initial branch exhibited a bootstrap value of 96%, indicating 96% similarity to the immediate ancestor. The initial progenitor possessed two daughters, MT874782.1and MT874783.1, which were unbranched and originated from a cluster of branches that were 96% genetically similar to the ancestral progenitor. Furthermore, these two daughters had normal TET2 samples. Within this cluster, the majority of TET2 sequences (23), normal controls (10), and AML patients (13) were found. This is interesting because it shows that TET2 sequences in clinical samples possess identical genetic characteristics. The normal and AML samples related to the ancestral healthy origin and the (13) AML samples at the end of the phylogenetic tree showed a huge mutation (deletion) that occurred in the AML samples, as shown in detail in (Fig.1c) and (Fig.2). In addition, the neighbor-joining tree (Fig.3) with bootstrap support values demonstrated a clear separation between most AML samples and normal controls, indicating that the mutations observed in AML samples represented significant evolutionary departures from the normal control TET2 sequence.

The 19^th^ branches developed from the 1st branch had bootstrap values of 85%, 80%, 80%, 57%, 61%, 58%, 100%, 100%, 62%, 61%, 61%, 61%,61%, 62%, 62%, 62%, 62%, 62%, and 62%, respectively, indicating the percentage of genetic relatedness to their progenitors. These bootstrap values provided statistical support for branching, indicating the reliability of these evolutionary relationships. These branches contain (10) normal controls, TET2 samples, and (13) AML patient samples, depicting the evolution over time. From the ancestral tree branches to subsequent generations of AML patients, there was a noticeable pattern of increased genetic diversity among the samples, indicating more evolutionary differences than similarities.

Notably, (10) normal control sequences clustered tightly near the ancestral root, reflecting a conserved function. Conversely, (13) AML samples formed a terminal cluster with long branches, suggesting a common mutational signature or evolutionary trajectory among AML patients, signalling accelerated divergence, which is a hallmark of clonal evolution[20].

Strikingly, the 9^th^ branch developed from the 8^th^ branch and further developed into the 10^th^ branch, depicting its evolution. These branches represented 100%, 100%, and 62% of their corresponding ancestors, respectively. Throughout the branches of the phylogenetic tree, from ancestors to descendants, there was a pattern of greater genetic diversity among samples, resulting in more evolutionary differences than similarities.

Each branch originating from the parent branch within the AML patient cluster had an identical percentage of similarity to its immediate parent, suggesting a consistent or highly similar pattern of TET2 effects in AML patients. Diversity over time may translate to the evolution of clinically relevant attributes of TET2 sequences, such as resistance to therapy and the capacity to transmit this resistance either laterally or vertically due to the acquisition of certain genetic elements.

### 3.4 Distance Matrix

In this study, a distance matrix was used to extensively compare sequences based on percentage identity and differences (nucleotide differences). The distance matrix serves as an indicator of the evolutionary relationships among samples, illustrating their proximity. The sequences were compared, and the relative distance between pairs was represented using numerical data and heatmap visualization. As the degree of relatedness increases, the value approaches a level greater than zero, indicating that individuals share the same genetic makeup with one another at a 100% identity level. The Percentage Identity matrix (Fig.4A) displays the percentage similarity between all the sequence pairs in the dataset. When the gap between the two aligned pairs widened, the percentage of identity decreased. A diagonal line passing through the rectangle to divide it into two identical triangles, indicates that there was no gap between the two aligned sequences. (Fig.4A) shows high intra-group identity (95-100%) within normal samples, high intra-group identity (98-100%) within AML, and lower inter-group identity (85-95%) between normal and AML samples. The clear pattern of higher identity within groups than between groups statistically supports the clustering observed in phylogenetic analyses.

**Figure 4A:**
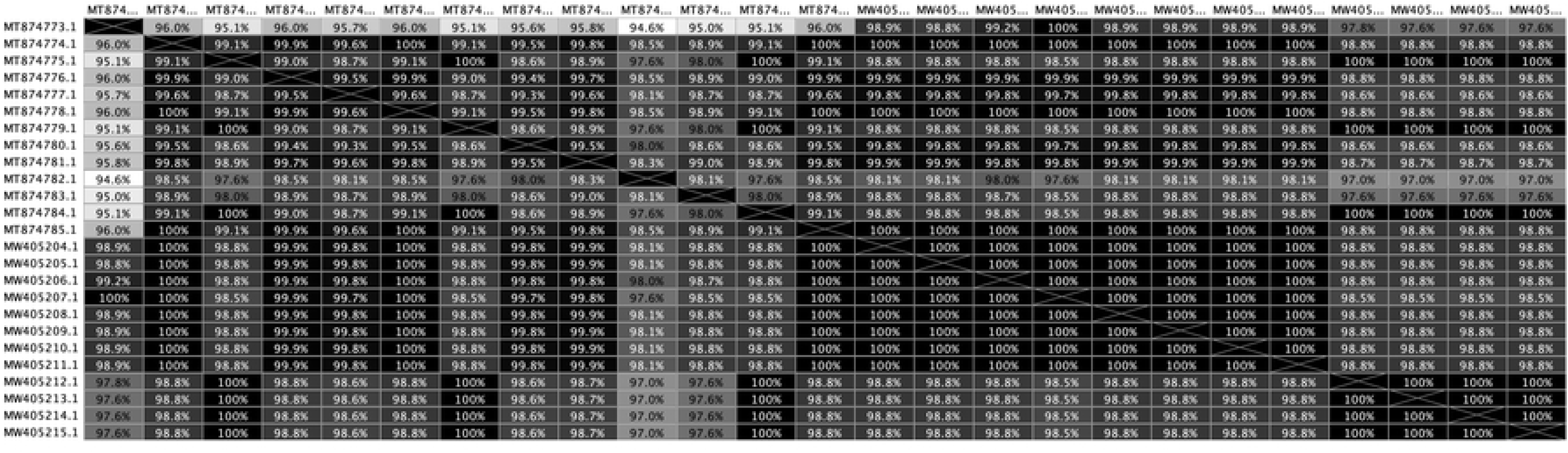
Percentage Identity (Distance Matrix / Heatmap) derived from the TET2 gene sequences related to AML patients and normal from Pakistan .100% indicates highly significant sequence correlation. The diagonal line traversing the rectangle, which bisects it into two triangles, signifies the absence of any distance between the two aligned sequences.

The total differences in nucleotide bases (Fig.4B) indicate the absolute number of nucleotide differences between sequence pairs. For example, comparing MT874782.1 (a sequence from normal samples) and MT874773.1 (a sequence from normal samples) showed 65 total differences in nucleotide bases, while MW405214.1 (a sequence from AML patient sample) and MW405204.1 (a sequence from AML patient sample) have (8) total differences in nucleotide bases. Furthermore, MW405214.1 (a sequence from AML patient sample) and MT874782.1 (a sequence from normal sample) had 34 total differences in nucleotides. Numerical differences directly quantify the genetic divergence that drives phylogenetic separation. Despite the sequences beingn from the same country, these results indicate a high rate of genetic diversity between samples from AML patients and those from normal patients. In other words, diversity is expected, which signifying a high rate of genetic diversity.

**Figure 4B:**
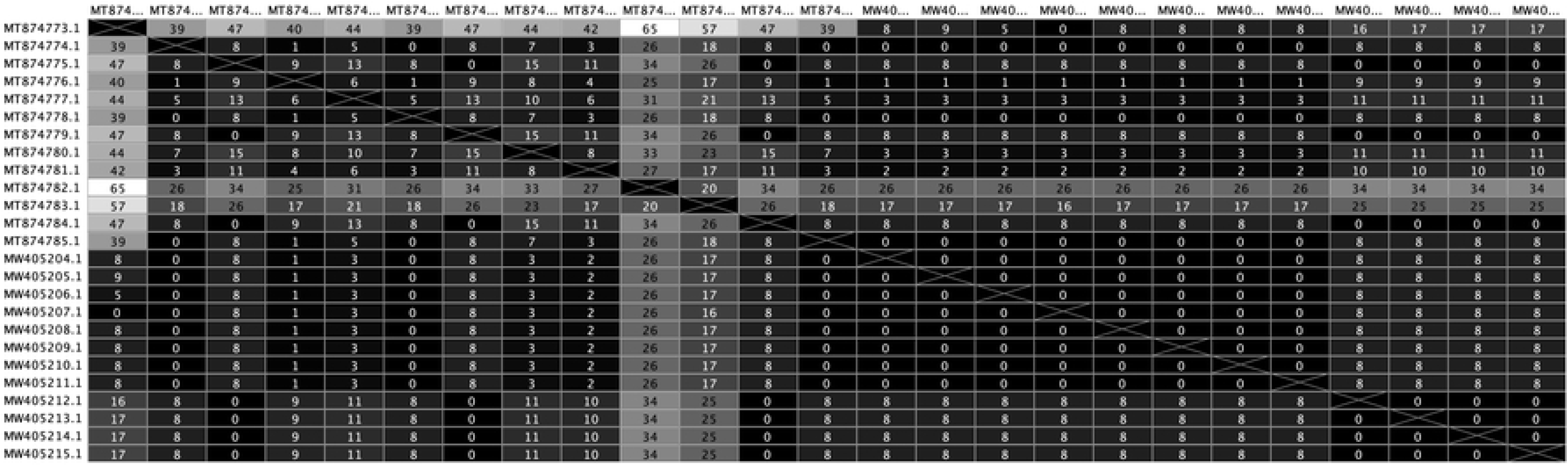
Total Differences in Nucleotide Bases (distance matrix/heatmap) derived from the TET2 gene sequences related to AML patients and normal individuals from Pakistan.The diagonal line traversing the rectangle, which bisects it into two triangles, signifies the absence of any distance between the two aligned sequences

Distance matrix analysis revealed different degrees of genetic deviation between the samples, whereby AML samples showed a greater degree of inter-sample variation than normal controls. This higher rate of variability in AML samples is in line with the clonal evolution theory of cancer, which suggests that disease progression and the development of therapeutic resistance are driven by the accumulation of mutations.

### 3.5 Population-Specific Patterns and Clinical Implications

Analysis of Pakistani TET2 sequences has revealed population-specific mutation patterns that may have implications for disease susceptibility and therapeutic response. The high frequency of deletions in the catalytic domain (positions 693-876) in AML samples suggests that this region is particularly vulnerable to mutagenic events in this population.

These findings align with recent studies indicating that TET2 mutations are prevalent in cytogenetically defined intermediate-risk AML, with a mutation frequency ranging from 18% to 23%[19]. The population-specific patterns observed in our study may contribute to the understanding of why certain populations have different susceptibilities to AML or different responses to therapeutic interventions.

### 3.6 Therapeutic Implications

Identification of particular mutational hotspots within TET2 has important therapeutic implications. Therefore, mutations in TET2 can be used to predict response rates to hypomethylating agents in patients with T-cell acute lymphoblastic leukemia (T-ALL), myelodysplastic syndrome (MDS), and AML, as these mutations sensitize cells to therapy[1], [19], [30], [36].

Comprehensive deletion of the TET2 catalytic domain in AML suggests the consideration of inhibitors of DNA methyltransferases and other epigenetic agents in AML patients. Identifying mutational patterns in different populations could result in a more personalized treatment method.

### 3.7 Comparison with Recent Literature

Our results align with those of the current comprehensive review of TET2 mutations in AML patients. In 2024, a study showed TET2 mutations in 7–28% of adult patients with AML, which were correlated with higher white blood cell counts and abnormal clinical outcomes[1].The significance of TET2 mutations in patients with AML has limited prognostic implications, with most studies proposing no significant effect on event-free survival, relapse rates, or overall survival between patients with and without TET2 mutations[1], [11].

Recent developments in single-cell genomics have indicated that cancer proliferation and clonal evolution are more complex than previously thought. Tools such as the SPRINTER algorithm, developed for single-cell whole-genome DNA sequencing data, have indicated that proliferation differs among evolutionarily distinct clones within identical tumors[51]. Further research is needed to examine single-cell approaches for analyzing clonal heterogeneity of TET2 mutations in AML patients.

### 3.8 Study Limitations and Future Directions

A number of limitations should be recognized with respect to our findings. The relatively small sample size (25 sequences) limited the statistical power for detecting population-specific patterns in this study. Furthermore, amplicon-based sequencing methods are unlikely to be comprehensive and may miss much information regarding clonal heterogeneity in AML samples. Recent single-cell sequencing methods have revealed that tumors contain numerous genetically distinct subpopulations, and more thorough investigations should use these methods[51], [52], [53].

Further studies should aim to increase the sample size to include larger cohorts, use single-cell sequencing technologies, and conduct functional studies to validate the biological relevance of identified mutations. In addition, longitudinal analysis of howTET2 mutations change over time in terms of disease progression may be an effective approach to learn more about the dynamics of alterations in TET2 mutations.

## 4. Conclusion

Phylogenetic analysis of the TET2 gene in Pakistani AML patients using sequence phylogeny and normal controls has provided substantial implications regarding the population-specific mutation patterns and evolutionary history of theTET2 gene involved in acute myeloid leukemia. Our study found that AML-derived TET2 sequences had substantial structural alterations, notably massive deletions in the catalytic domain (positions 693-876), which are essential for TET2 enzymatic activity and the demethylation pathway.

The population-specific characteristics of TET2 mutations may have important implications in determining disease susceptibility, evolution, and therapeutic response in the Pakistani population. The higher variability in GC content and sequence structure in AML samples compared to normal controls supports the role of TET2 mutations as the underlying cause of leukemogenesis by disrupting normal epigenetic regulation.

Our results are consistent with the rapidly growing evidence that TET2 mutations occur in 7-28% of adult AML patients and are associated with altered clinical outcomes. The clinical translation of the determination of specific mutational hotspots is the basis for reducing the current therapeutic profile towards the development of appropriate therapeutic plans aimed at the treatment of such hotspots, especially with hypomethylating agents and other epigenetic regulators.

However, multiple limitations must be considered, such as the relatively small sample size, and the use of amplicon sequencing technologies, which may underrepresent the extent of clonal heterogeneity. Future research should use single-cell sequencing technologies and increasingly diverse and larger cohorts to replicate and expand these results.

This study reinforces the growing understanding of population-specific genetic variation in cancer, and the necessity of considering an evolutionary perspective in the study of cancer genomes. The insights obtained in this study can be used to develop specific treatment regimens for AML patients, especially for populations with certain genetic backgrounds. In summary, the TET2-related sequence data analyzed here provides novel insights into the evolution of AML at the molecular level, identifying the pivotal role of TET2 structural modifications in the occurrence of the disease and possible directions for individualized treatment strategies.

It is important to recognize the limitations of this study. Primarily, the sample size, comprising 25 sequences (12 AML and 13 normal),was relatively small for robust phylogenetic analysis, which may limit the statistical power of our conclusions. Second, the use of amplicon sequencing approaches may not capture the clonal heterogeneity present within individual AML samples, as modern single-cell sequencing methods have shown that tumors contain multiple genetically distinct subpopulations[52], [53].Third, population-specific claims require validation with larger and more diverse cohorts to establish statistical significance. Future studies should consider employing single-cell whole-genome DNA sequencing (scDNA-seq) approaches, which have been shown to provide a more accurate identification of clonal evolution and proliferation dynamics in cancer[51].

## 5. Future Directions

Based on our findings and current developments in this field, several important directions should be pursued to deepen our understanding of TET2’s role in AML.

### 5.1 Single-Cell Genomics Approaches

Future research should use single-cell whole-genome DNA sequencing (scDNA-seq) technologies to gain a better understanding of the clonal heterogeneity of TET2 mutations in AML. Recent innovations, including the SPRINTER algorithm, have shown that proliferation differs across evolutionarily distinct clones within a single tumor, providing more accurate insights into cancer evolution than bulk sequencing methods[51].

### 5.2 Functional Validation Studies

Further functional studies are required to validate the biological relevance of the identified mutations, especially those that include large deletions in the catalytic domain. Such studies are needed to determine how certain mutations in TET2 are associated with altered TET2 enzymatic function, DNA methylation, and genes involved in the expression profiles.

### 5.3 Therapeutic Response Studies

Given that TET2 mutations may indicate responsiveness to hypomethylating agents, clinical studies should investigate the therapeutic efficacy of these agents in patients with the specific TET2 mutation patterns identified in this study.

### 5.4 Expanded Population Studies

Population-specific patterns need to be validated by examining larger cohorts, such as populations representing a wider age range or including a wide variety of AML subtypes. This would assist in determining whether the patterns present in the Pakistani population are unique or part of a larger regional or ethnic difference.

## Acknowledgments

We thank the National Center for Biotechnology Information (NCBI) for providing access to the sequence data used in this study. We are also grateful to the research community whose efforts have contributed to our Knowledge of TET2 biology and AML pathogenesis.

## Data availability statement

All sequence data used in this study are publicly accessible from the National Center for Biotechnology Information (NCBI) database (https://www.ncbi.nlm.nih.gov/).Table1 listed the accession numbers for all sequences.

